# Interindividual Diversity of Human Gut Mucin-Degrading Microbial Consortia

**DOI:** 10.1101/2023.05.13.540604

**Authors:** Ashwana D. Fricker, Tianming Yao, Stephen R. Lindemann, Gilberto E. Flores

## Abstract

Mucin is a glycoprotein secreted throughout the mammalian gastrointestinal tract that can support endogenous microorganisms in the absence of complex polysaccharides. While diverse mucin degrading bacteria have been identified, the individual host microbial community differences capable of metabolizing this complex polymer are not well described. To determine whether individuals have taxonomically distinct but functionally similar mucin-degrading communities, we used a ten-day *in vitro* sequential batch culture fermentation from three human donors with mucin as the sole carbon source. For each donor, 16S rRNA gene amplicon sequencing was used to characterize microbial community succession, and the short-chain fatty acid profile was determined from the final community. Although two of the final communities had genus-level taxonomic differences signified by the presence of *Desulfovibrio* and *Akkermansia*, other members, such as *Bacteroides*, were shared between all three final communities. Metabolic output differences were most notable for one of the donor’s communities, with significantly less production of acetate and propionate than the other two communities. These findings reinforce the concept of a taxonomically distinct and, at broad levels, a functionally redundant gut microbiome. Furthermore, the mechanisms and efficiencies of mucin degradation across individuals are important for understanding how this community-level process impacts human health.

## Introduction

Everyone has a personalized intestinal microbiome, which assists in normal digestion that is cultivated through their lifetime and tightly linked to both health and disease. The composition of each individual’s gut microbiome is highly dependent on nutrient intake (Mohammadkhah et al. 2018). Although individual microorganisms can be associated with some disease states, other diseases are associated with microbial community features, examples include organisms that are efficient at dietary energy extraction associated with obesity (B.-N. Liu et al. 2021) and organisms that contain sulfur metabolic machinery associated with colorectal cancer (Wolf et al. 2022). These microorganisms may affect host health through the production of metabolites that interact with local and distal tissues (Parada Venegas et al. 2019). For example, the production of short-chain fatty acids (SCFA) by fermentative gut bacteria has been implicated in protection of the epithelium, immune regulation, and epithelial cell proliferation and differentiation (Bilotta and Cong 2019). The mechanisms that maintain and bolster these functions of the microbiome are poorly understood but of great interest because of their potential to modulate health.

Microorganisms within the intestinal tract require basic resources including carbon and energy for growth and survival, which they derive primarily from the host diet (Leeming et al. 2019). Dietary fibers, as one example, compose a broad array of polysaccharides exclusively metabolized by diverse members of the gut microbiome. Recent evidence indicates that dietary fibers with more complex chemical structures necessitate a diverse array of enzymes for digestion often produced by specialist microorganisms (Lindemann 2020). These specialists enable a myriad of microbial interactions as they liberate smaller sugar subunits and produce fermentation products that are both used by other members of the microbiome and by host cells (Koropatkin, Cameron, and Martens 2012). Comparatively simple fibers require fewer enzymes for degradation, making these polysaccharides more widely accessible to many gut organisms (Cantu-Jungles et al. 2021; Romero Marcia et al. 2021). Therefore, variability in host diets leads to a difference in dietary fibers consumed, ultimately conferring advantages to specific sets of microorganisms capable of breaking the discrete bond types and helping to shape the uniqueness of each microbiome.

Despite this individuality in diet, health humans secrete mucins, a family of glycoproteins decorated with O-linked carbohydrates in various configurations produced by specific epithelial cells in the intestinal, respiratory, and genitourinary tracts. Although primarily known for their role as a barrier against invasion, the high polysaccharide content of mucins can potentially serve as a rich and reliable nutrient source (Tailford et al. 2015). Mucin structure is complex, fundamentally containing a serine-threonine-proline rich protein backbone, an N-acetylgalactosamine (GalNAc) O-linked to this protein, an array of sugars linked to the GalNAc, and a final terminal unit (Corfield 2015). The breadth in structures lies in the sugar array, which can comprise α-(1-3)- or β-(1-3)-linked galactose, or a β-(1-6)-, β-(1-3)-, α-(1-6)-linked N-acetylglucosamine (GlcNAc). Furthermore, these structures can be capped by a sulfate group, fucose, or N-acetylneuraminic acid (sialic acid; NeuAc). Given this diversity of chemical linkages, only a subset of gut microorganisms possess the functional capacity to hydrolyze and utilize mucins for growth (Glover, Ticer, and Engevik 2022).

Some organisms, such as the nutritional generalist, *Bacteroides thetaiotaomicron*, can switch between fermentation of dietary polysaccharides and mucin glycans (Ravcheev et al. 2013). During fermentation of either dietary fibers or mucin, byproducts, including the SCFA acetate, are produced (Mahowald et al. 2009). These metabolites stimulate mucin production by goblet cells in the gut, resulting in an intact mucus barrier between human and bacterial cells along the intestines (Willemsen et al. 2003; Adamberg et al. 2018). In fact, there appears to be a direct connection between the abundance of dietary fiber in the diet and colonic mucin thickness (Brownlee et al. 2003; Hedemann, Theil, and Bach Knudsen 2009; Earle et al. 2015; Desai et al. 2016), likely mediated by the microbiota.

A smaller subset of organisms in the gastrointestinal tract, like the mucin-degrading specialist *Akkermansia muciniphil*a, appear to depend on direct mucin fermentation for growth (Derrien et al. 2004). There are many carbohydrate active enzymes (CAZymes) responsible for metabolizing complex mucin glycans, where each enzyme (i.e. glycoside hydrolase – GH) is responsible for hydrolysis of a specific glycosidic bond, liberating specific sugar moieties. Although it is possible that *Akkermansia*, a specialist that retains a full suite of mucin-degrading CAZymes, is capable of filling this mucin-degrading niche and outcompeting other organisms for this resource, the incidence rate is only 50-80% of the human population (Geerlings et al. 2018). Given that much of the established abundances of microorganisms are confined by available technology, including extraction methods, primer choice, and detection limits (Brandt and Albertsen 2018; Abellan-Schneyder et al. 2021), *Akkermansia* may in fact be present in all gut communities. Alternatively, it is possible that gut microbial communities divide the metabolic labor in which partial hydrolysis of mucin releases mono- or di-saccharides that can then be fermented by other organisms (Bunesova, Lacroix, and Schwab 2018). In this second scenario, the activities of secondary consumer microbiota would be supported by the intestinal mucin and further stimulate mucin production, thereby limiting pathogen invasion. To determine whether mucin metabolism would be monopolized by a single member or support a complex community, we sought to characterize the indigenous microbial communities capable of growth on mucin across multiple individuals.

Given the prevalence in known mucin degraders, we hypothesized that the communities capable of metabolizing mucin between two individuals are likely to be taxonomically distinct, have a set of functional redundancies with respect to mucin degradation, and produce SCFA profiles that depend on the functional capacity of the community. Here, using a top-down approach to select for mucin-degrading communities from three individuals, we identify a core and unique set of microorganisms involved in mucin metabolism with distinct short-chain fatty acid production profiles.

## Methods

### Medium

Fermentation Buffer was based on Yao and colleagues (Yao, Chen, and Lindemann 2020), with modifications. Briefly, our fermentation buffer contained 24 μM Na_2_HPO_4_, 15.6 μM NaH_2_PO_4_, 8 mM NaCl, 6 mM KCl, 6.7 mM NH_4_Cl, 0.7 mM Na_2_SO_4_, 1 ug/mL Resazurin, 3.2 mM Urea, 1.4 mM Cysteine-HCl, 0.5 mM CaCl_2_-2H_2_O, 0.5 mM MgCl_2_-6H_2_O and trace minerals solution. Trace minerals solution is based on Ferguson and Mah (Ferguson and Mah 1983), where the final composition in fermentation buffer consisted of 1.3 μM Na_2_-EDTA -2H_2_O, 0.6 μM CoCl_2_ - 6H_2_O, 0.5 μM MnCl_2_ -4H_2_O, 0.36 μM FeSO_4_ -7H_2_O, 0.7 μM ZnCl_2_, 0.17 μM AlCl_3_ -6H_2_O, 0.09 μM Na_2_WO_4_ -2H_2_O, 0.12 μM CuCl_2_ -2H_2_O, 0.076 μM NiSO_4_ -6H_2_O, 0.077 μM H_2_SeO_3_, 0.16 μM H_3_BO_3_, 0.04 μM NaMoO_4_ -2H_2_O. To alleviate amino acid auxotrophies, either 10 μM equimolar amino acid mix or 0.1 mg/ mL tryptone (Oxoid) was added. Tryptone concentration was normalized to the amino acid mix based on the percentage of L-cysteine as published by USBiological Life Sciences (“Tryptone CAS 91079-40-2” n.d.).

#### Soluble Mucin Preparation

Preparation of soluble mucin was based on Kirmiz and colleagues (Kirmiz et al. 2020) and added to a final concentration of 0.5%. Briefly, soluble porcine gastric mucin type III (Sigma Aldrich, USA) was prepared by autoclaving a 5% (w/v) solution in 0.01 M phosphate buffer (6.5 mM KH_2_PO_4_ and 3.5 mM K_2_HPO_4_), dialyzing with a 12- to 14-kDa membrane (Spectra/Por 4; Spectrum Laboratories, Rancho Dominguez, CA) overnight in 10 dialysis volumes of 0.01M phosphate buffer, centrifuging for 30 min at 30,000Lrcf, and filtering through a 0.45-μm syringe filter followed by a 0.2-μm syringe filter (Whatman GE Healthcare Life Sciences, Chicago, IL). Sterile soluble mucin was then added to the fermentation buffer at a final concentration of 0.5%.

#### Mixing of mucin solution and fermentation buffer

Briefly, concentrated fermentation buffer (2X) was aerobically prepared and diluted to a working concentration with soluble mucin and sterile water. Prior to transfer to Balch tubes, buffer-mucin mix was stored covered in the anaerobic chamber (85% N_2_, 10% CO_2_, 5% H_2_) for 48h to degas. Each tube received 5 mL fermentation buffer except the initial three tubes which each received 4 mL fermentation buffer. Fermentation buffer containing mucin was aseptically distributed into Balch tubes in a 10% CO_2_, 5% H_2_, and 85% N_2_ atmosphere before sealing with butyl rubber stoppers. After preparation, tubes were stored at 37°C prior to inoculation to detect any contamination. Immediately before inoculation, 0.05 mL ATCC vitamin mix [final 1% v/v, manufacturer’s number: ATCC MD-VS, Hampton, NH] was added to each tube. An additional three tubes were maintained for the full duration of the experiment at 37°C without inoculation as negative controls.

### Inoculum Preparation

Approximately 4 ± 1.5 g of fecal material was collected from three healthy donors. Donors (ages 20-45 yrs) had not received any antibiotic treatment within three months and were generally in good health. The protocol for collecting human feces was approved by the Institutional Review Board of California State University, Northridge (#1516-146-f). Donors were asked to collect two Hershey’s kiss size samples (∼4.2 g each) into separate pre-weighed conical vials using sterile wooden spatulas. Samples were maintained on ice during transfer and received within three hours of evacuation. Fecal material was diluted 1:4 w/v with 1X fermentation buffer containing mucin as described above. Tubes were vortexed for 20 seconds, returned to ice for up to 5 minutes, and vortexed again for 20 seconds. The fecal slurry was then poured over 4-ply sterile cheesecloth, and the flow through was collected with a syringe. Each tube containing 4 mL of fermentation buffer received 1 mL of fecal slurry totaling a 1:20 fecal dilution. The initial inoculation tubes received an additional 50 μL of 0.25 g/mL cysteine to reduce the initial inoculum for a final total cysteine concentration of 2.4 mM. Tubes were placed in a static 37°C incubator inside of an anaerobic Coy chamber (10% CO_2_, 5% H_2_, and 85% N_2_ atmosphere) for 23 to 25 hours before sample collection and transfer as described below.

### In vitro sequential fecal fermentation

The sequential cultivation experiment was continued for 10 consecutive days, and each donor was cultured in triplicate lineages that were not intermixed (Yao, Chen, and Lindemann 2020). Daily, immediately prior to inoculation, 50 μL of ATCC vitamins (final 1% v/v, manufacturer’s number: ATCC MD-VS, Hampton, NH) was added to each 5 mL fermentation buffer. Due to settling of microorganisms from static incubation, tubes were gently inverted 5x prior to sampling. Subsequently, 50 μL of the incubated culture was transferred to the corresponding tube containing 5mL fermentation buffer and incubated in the Coy chamber as described above.

Daily, 500 μL of each incubated culture was collected and centrifuged at 10,000 x g for 5 minutes. The cell pellet was stored at −20°C until thawing for DNA extraction. On the final day, after centrifugation, the supernatant was transferred to a separate microfuge tube and stored at −20°C until thawing for SCFA detection as indicated below.

### DNA Extraction and Sequencing

Pellets were extracted following the Zymo DNA Microprep Kit #D4301 (Zymo Research Corporation, USA), with the following modifications. For the initial inoculum, 20% of the resuspended culture was processed to prevent overloading the kit. After resuspension, samples were placed on a horizontal vortex adapter at max speed for 10 minutes. DNA concentrations were read on a Qubit 2.0 fluorometer with the high-sensitivity kit, following manufacturer’s protocol. One microliter of each eluted sample was used in polymerase chain reaction amplification of the variable region 4 of bacterial and archaeal 16S ribosomal RNA genes with barcoding primer set 515/806 based on the original Earth Microbiome Project protocol (Caporaso et al. 2011; “16S Illumina Amplicon ProtocolL: Earthmicrobiome” n.d.). Primers used were 515F (Parada, Needham, and Fuhrman 2016) - GTGYCAGCMGCCGCGGTAA) and (806R - GGACTACHVGGGTWTCTAAT) with barcoding on the 806R primer. PCR mixture conditions and thermal cycling steps are previously described (Herman et al. 2020). Triplicate PCR reactions for each sample were combined, quantified with Quant-iT PicoGreen (Invitrogen) on a SpectraMax3 [Molecular Devices, USA], and pooled at equimolar concentrations before single-tube cleaning with the Invitrogen Gel and PCR Clean-up Kit (Invitrogen) and sequencing using an Illumina MiSeq (v2, 2 × 150 bp) instrument (Illumina, USA).

### DNA Sequence Analysis

Sequences were analyzed following the QIIME-2 Atacama Desert pipeline (Qiime2Docs), briefly described here. Sequences were denoised, dereplicated, and chimeras were removed with Dada-2 (Callahan et al. 2016) with the following parameters: --p-trim-left 5 - -p-trunc-len 150. Amplicon sequence variants (ASVs) were classified using sklearn and the silva 138.99 database (Quast et al. 2012). PCR blanks were used to remove contaminating ASVs following the decontamR pipeline (Davis et al. 2018) using a prevalence-based strategy. QIIME-2 was used to calculate various α-diversity (total ASV counts and Shannon Diversity Index) and β-diversity (Bray Curtis) metrics.

### Determination of Fatty Acids

Final consortia cultures at day 10 were analyzed for SCFAs of acetate, propionate, butyrate and BCFAs of isobutyrate and isovalerate. Samples were measured on a GC-FID system (GC-2030, Nexis, Shimadzu Corporation, Kyoto, Japan). Cultures were centrifuged (10L000 × g) for 5 minutes to remove cell debris, mixed 4:1 with an internal standard mixture containing 4-methylvaleric acid, phosphoric acid, and copper sulfate pentahydrate, and supernatants were transferred to 2 mL screw-thread autosampler vials. Volatile compounds were quantified on a GC-FID with a wax column (RESTEK Stabilwax-DA #11023, Bellefonte, PA). Helium was used as a carrier gas for the column at a flow of 1.41 mL/min. Samples were injected in a split mode (5 uL injection volume, 1:5 split ratio) under a linear velocity of 35 cm/s at 230 °C. The oven temperature was set as below: initial temperature at 40 °C for 2 min, following a ramp to 130 °C in 3 min, and then another ramp to 195 °C in 6.5 min, holding at 195 °C for 4 min.

### pH and OD Testing

On the final day of sampling, tubes were removed from the anaerobic chamber and opened aerobically. Cultures were transferred to a 15 mL conical vial and 100 μL of the culture was used to determine OD_600nm_ on a Biophotometer plus spectrophotometer (Eppendorf, Germany). Cultures with an OD_600nm_ > 1.0 were diluted with sterile water and re-read. The pH was tested on the remaining volume using an accumet pH meter (Fisher Scientific, USA).

### Statistical Analysis

For pH, OD_600nm_, SCFA, and alpha diversity analysis, statistically significant differences were calculated by Tukey’s honest significance multiple comparisons test using R 4.1.1 (R Foundation for Statistical Computing, Vienna, Austria). Comparisons were considered significant if corrected P-values were less than 0.05. For beta diversity analysis, dissimilarity matrices generated in Qiime-2 were used to calculate statistically significant differences with ADONIS from the vegan package in R 4.1.1. Comparisons were considered significant if the p value of the F statistic was less than 0.001. Symbol style for figures: nonsignificant (ns), 0.05 (*), 0.01 (**), 0.001(***) and <0.0001(****).

## Results

To better understand mucin degradation as a community level process in the human gut microbiome, we performed 10-day sequential fecal enrichments from three human donors in triplicate, using purified porcine gastric mucin as the sole carbon and energy source. As a secondary variable, we used two amino acid sources during community development in parallel enrichments for each donor. For each day, cultures were transferred at a 10^-2^ dilution rate and cell pellets were collected to monitor community development using 16S rRNA gene amplicon sequencing. On the final day, the OD_600nm_ and pH of cultures was tested, and the spent culture supernatant was collected and used to determine short-chain fatty acid production.

### Amino acid source does not govern growth or metabolic outputs

In cultures from all three donors, robust growth was evident by changes in media turbidity after each day of incubation, regardless of amino acid treatment. On the final day of growth, all communities grew to OD_600nm_ > 1 and there were no significant differences in OD_600nm_ within a donor’s communities on either amino acid source (Supplemental Figure 1). However, across donors, communities from donors one and three grew to significantly higher OD_600nm_ (∼1.4-1.6) than donor two (∼1.1) suggesting difference in growth yield (Figure 1A). Similarly, drops in pH were also generally greater for communities from donors one and three, except for on the amino acid mixture where donor two and three communities both dropped ∼0.3 pH units (Figure 1B). Interestingly, communities from donor two had a significantly different drop in pH between the two amino acid sources, with the culture growing on mixed amino acids dropping to pH 6.1 and the culture grown on tryptone to 6.3 (Supplemental Figure 1). This likely suggests ammonification of the medium from degradation of the more abundant proteins in the community cultivated on tryptone.

**Figure 1.**
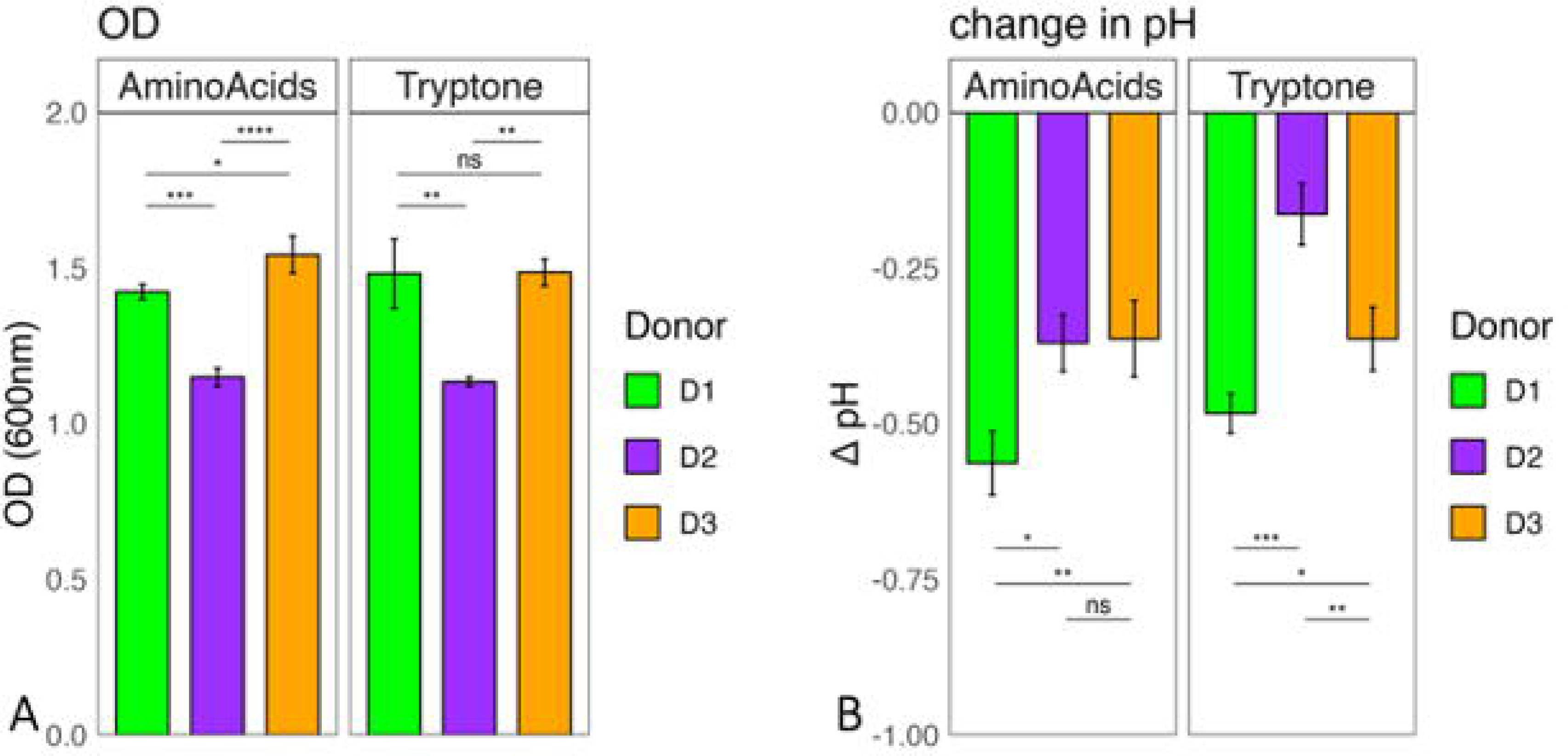
Mucin degrading consortia from three human donors established on different amino acid sources grew to different final optical densities (A) and produced different levels of acid (B) after 10 days of incubation. Averages of three lineages from each final donor community ((Donor 1, D1; Donor2, D2; Donor3, D3) are displayed with error bars representing standard deviation as calculated in R. Statistically significant differences are calculated by Tukey’s multiple comparisons test with P<0.05. Symbol style: nonsignificant (ns), 0.05 (*), 0.01 (**), 0.001(***) and <0.0001(****)

### Short-chain fatty acid output is unique to each community

SCFAs are main metabolic byproducts of gut microbial fermentation of complex carbohydrates. Therefore, to determine the metabolic output of each community during mucin fermentation, the SCFA profile was assayed on day 10 of sample collection (Figure 2). Overall, microbial consortia from donors one and three produced the highest total concentration of SCFAs (∼20 mM, sum of acetate, butyrate, and propionate) whereas the donor two community produced lower concentrations (14 mM), irrespective of amino acid source. These differences were largely driven by acetate and propionate, where cultures from donors one and three produced more than donor two. However, differences of specific SCFAs generated by each donor’s communities were dependent on amino acid source as discussed below.

**Figure 2.**
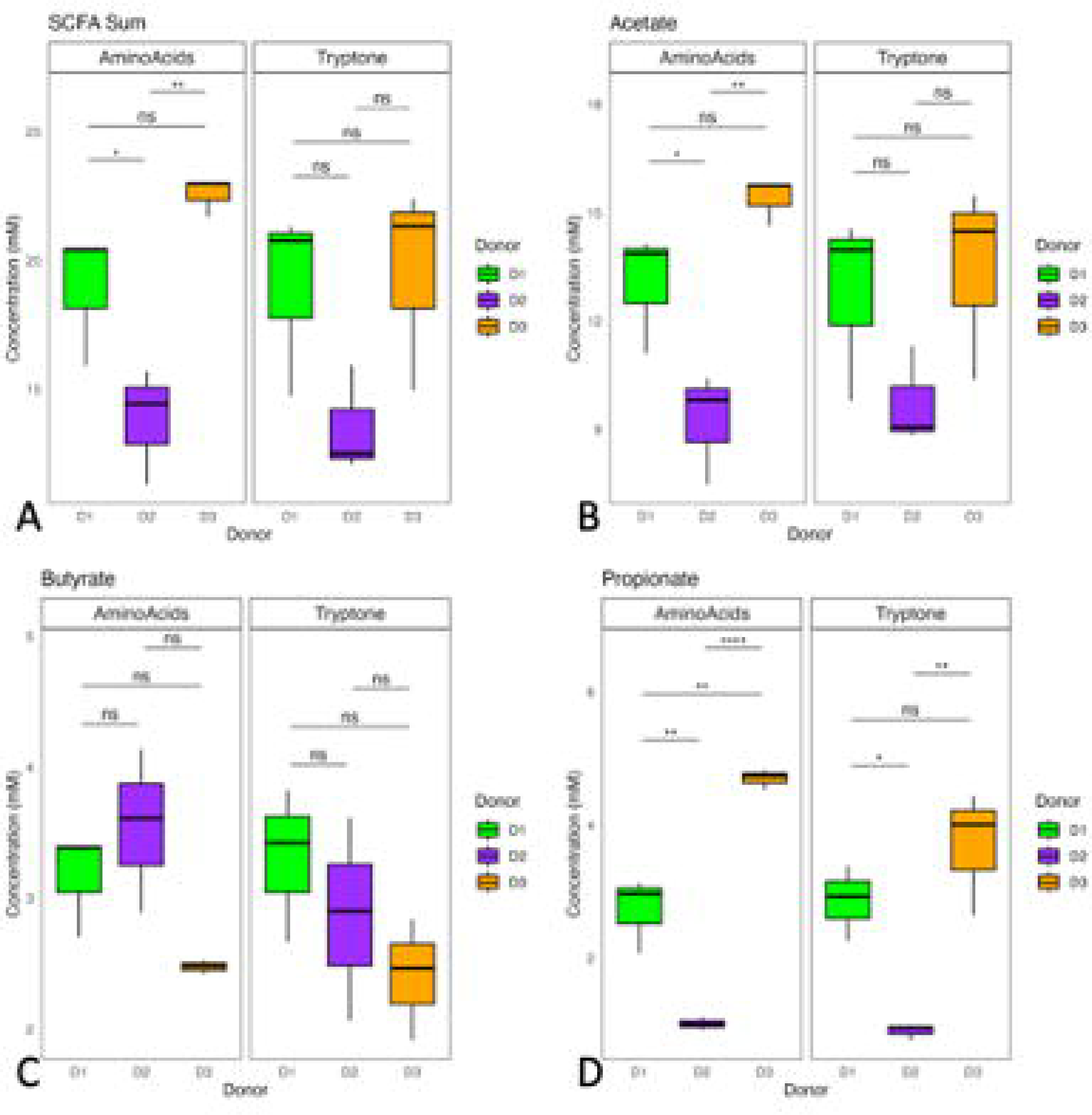
Mucin degrading consortia from donors one and three produced the highest total concentration of SCFAs driven by acetate and propionate, whereas the donor two consortium produced lower concentrations, irrespective of amino acid source. Mean, first and third quartiles of total SCFAs (A), acetate (B), propionate (C), and butyrate (D) for each final donor community (Donor 1, D1; Donor2, D2; Donor3, D3) are represented. Statistically significant differences are calculated by Tukey’s multiple comparisons test. Symbol style: nonsignificant (ns), 0.05 (*), 0.01 (**), 0.001(***) and <0.0001(****)

When cultivated on tryptone, communities from donors one and three produced equal amounts of propionate (3-4 mM), whereas the donor two consortium produced significantly less (0.5 mM). In contrast, there was no significant difference in the production of acetate (9-14 mM) and butyrate (2.5 – 3.5 mM) across the three communities. Although the same general trend of SCFA production was observed in cultures grown with the amino acid mix, the differences between communities was more pronounced. In these communities, donor three microbiota produced the greatest amounts of propionate (5 mM), followed by donor one microbiota (2.5 mM), with donor two’s microbiota generating low concentrations (0.5 mM). Likewise, concentrations of acetate produced by both donor one and donor three communities were similar (13-15 mM), with donor two’s producing lower concentrations (9 mM). The production of butyrate in amino acid mix-consuming communities mirrored those cultivated on tryptone, where there was no significant difference in the amount of butyrate produced (2.5 – 3.5 mM). Furthermore, we detected trace amounts of the branch chain fatty acids, isovalerate and isobutyrate (Supplemental Figure 2). These minor BCFAs reflected the patterns observed for SCFAs, but had very low concentrations. Overall, these data coupled with the OD_600nm_ results, suggest greater overall utilization of mucin by microorganisms from donors one and three.

### 16SrRNA amplicon sequence quality and taxonomy

Across two MiSeq Illumina sequencing runs, a total of 13,120,305 sequences were obtained. Each run was independently denoised, dereplicated, and chimeras were removed. Amplification blanks were used to remove contaminating sequences, where 66 ASVs from the first sequencing run and 9 ASVs from the second were removed. Given that the experiment occurred with triplicate lineages, after combining the two datasets, ASVs that were present in less than two samples were removed. Subsequently, samples with fewer than 5,000 reads, representing the extraction blanks, were removed. The final number of sequences analyzed constituted 92.9% of the original sequence total (12,184,363 sequences). To determine whether sequencing run bias would influence the results, six samples representing the initial inoculum and mid-to-end timepoints from both donors one and two were sequenced on both sequencing runs and community composition was compared (Supplemental Table 1). The differences in β-diversity (Bray-Curtis dissimilarity) across samples sequenced on the first and second run were not significantly different as determined by adonis analysis (R^2^=0.003, p = 0.95).

### Final community membership is dependent on initial inoculum

Six final enrichment cultures were developed from three donors to test whether the same final communities resulted in response to the high dilution and common carbon source. To determine community composition across time, the V4 region of the 16S rRNA gene was sequenced and relative abundances of the top thirty ASVs plotted. Across all donors, microbial succession was similar across the three replicate lineages from each donor in both basal buffers, suggesting that community composition is replicable under high dilution pressure with mucin. These data suggest that mucin structure deterministically governs communities, as has been observed for other complex carbohydrates (Yao, Chen, and Lindemann 2020). Although the compositions of mucin cultures on the first day of incubation cultures were similar to that of the initial fecal inoculum, the community increasingly diverged with each round of transfer. At day 5, the species richness plateaued, as determined by alpha diversity metrics including ASV count (Figure 3). There were no significant differences between the two amino acid sources for any of the three donors, whereas the initial diversity of the host microbiota appeared to determine the final diversity. Each donor started and ended with a different final number of ASVs, where donors one and two started with the greatest diversity at 162-179 ASVs and 157-170 ASVs, respectively. These communities plateaued to communities with 62 (donor one) and 46 (donor two) ASVs. Interestingly, donor three had much lower initial diversity, starting at about half the initial number of ASVs (91-101) compared to donors one and two, and plateaued to a correspondingly low number of 30 ASVs. However, the percent loss was approximately equal across donor samples, where each final community represented approximately 30% of the initial community (Donor 1, 35.5%; Donor 2, 29.7%; Donor 3, 31.8% Supplemental Figure 3). Furthermore, the point at which the loss in ASVs plateaued differed between the communities; communities belonging to donors one and two took 5 and 6 days, respectively, to reach this plateau, whereas donor three communities took 4 days. With the Shannon diversity metric, which accounts for both richness and evenness, we note the same trend in community diversity across three donors, where donor three communities leveled off faster than donors one and two.

**Figure 3.**
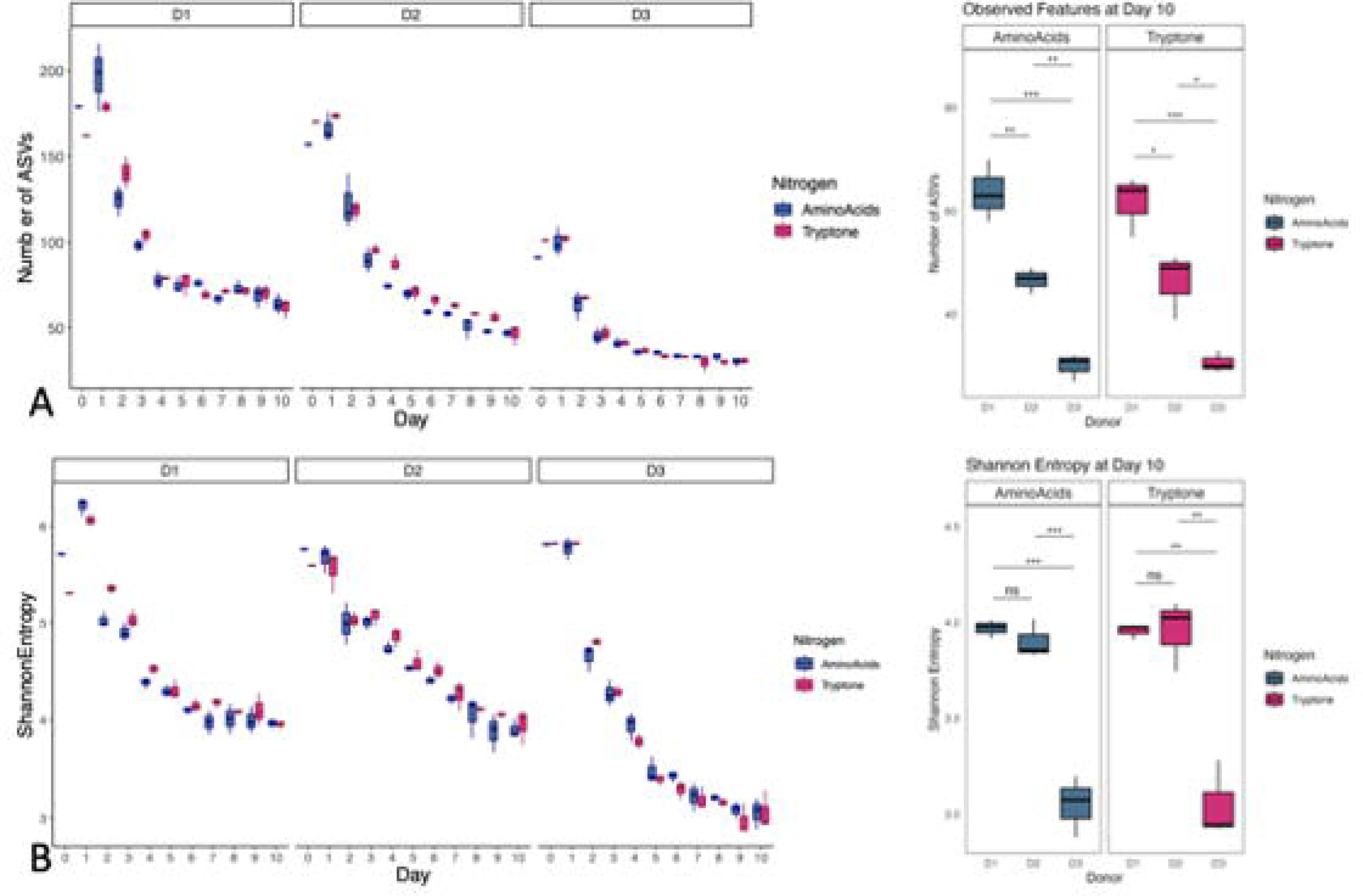
Species richness and evenness plateaued faster and reached lower final values for donor three communities as compared to donors one and two. Community membership at each passage for all three donors (Donor 1, D1; Donor2, D2; Donor3, D3) when supplemented with amino acids provided as an equimolar mix (Amino Acids, blue) or peptides (Tryptone, pink). Mean, first and third quartiles of the number of ASVs (A) and Shannon Entropy (B) are represented. Statistically significant differences at day 10 are calculated by Tukey’s multiple comparisons test. Symbol style: nonsignificant (ns), 0.05 (*), 0.01 (**), 0.001(***) and <0.0001(****)

Dominant phyla across all three communities included Bacteroidetes (Bacteroidota), Firmicutes (Bacillota), and Proteobacteria (Pseudomonadota) (Figure 4, Supplemental Figure 4). While there were no shared ASVs within the Bacteroidetes, by day 10 all three donor communities shared ASVs belonging to Firmicutes and Proteobacteria. At the genus level, these included organisms belonging to the Clostridium_innocuum group, *Hungatella*, *Lachnoclostridium*, *Faecalibacterium*, *Flavonifractor*, and *Escherichia-Shigella*.

**Figure 4.**
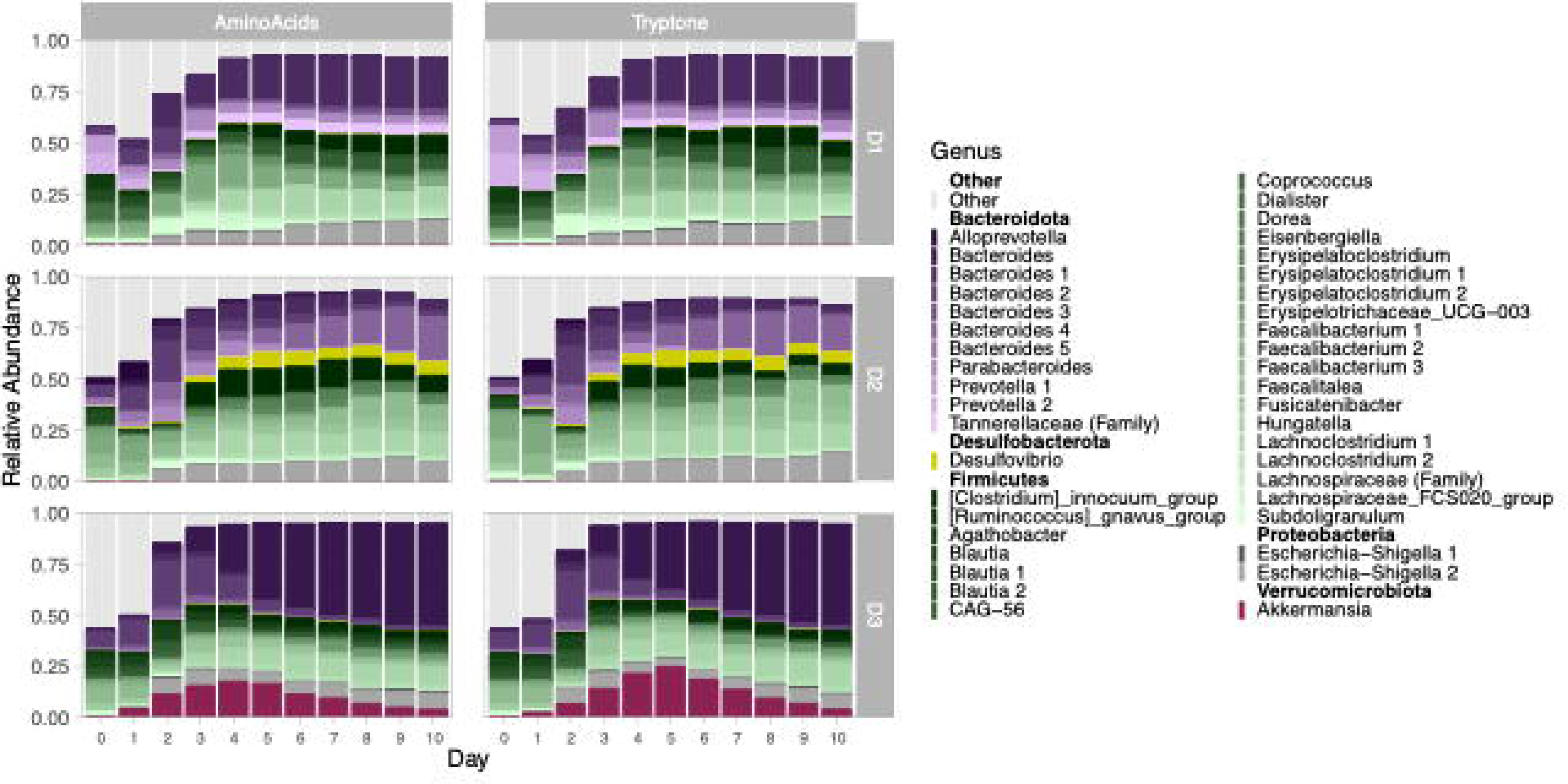
Mucin sustains diverse fermenting consortia over sequential dilutions that is dependent on the initial community. Averaged (n=3) relative abundance of community membership for each donor over 10 days with supplementation of either Amino Acids or Tryptone are shown. Each shade represents a distinct ASV, with the color representing a phylum: Bacteroidota (purple), Desulfobacterota (yellow), Bacillota (green), Proteobacteria (grey), and Verrucomicrobiota (red). The top fifty taxa across all donors (Donor 1, D1; Donor2, D2; Donor3, D3) are shown, where remaining low-abundance ASVs are grouped in Other (light grey).

In addition to shared features of the three communities, each also had unique membership. In donor two communities, a bloom of *Desulfobacterota* appeared at day three and remained abundant throughout the experiment. On the other hand, donor three displayed a bloom of *Akkermansia* that had the highest abundances on days 4 and 5, but whose population declined until the termination of the experiment on day 10. To identify less-obvious associations of individual taxa with their donors, we compared the taxa of the day 10 microbial communities using linear discriminant analysis effect size (LEfSe, Supplemental Figure 5). Interestingly, some of the organisms that were shared across the three donors were discriminants for a specific donor due to the differential abundance patterns. At the genus level, in addition to *Desulfovibrio,* other discriminating taxa included *Faecalitalea*, *Merdibacter, and Erysipelatoclostridium* for the donor two consortium. *Akkermansia* was a discriminant of the donor three community, as were *Ruminococcus* and *Oscillospiraceae*. Although donor one did not have any obvious associations with specific taxa in the visualizations, the FCS020_group and *Howardella* (in the Lachnospiraceae family), *Prevotella*, and *Megamonas* were all discriminants of this donor. Interestingly, at the ASV level, each donor had multiple ASVs belonging to the genus *Bacteroides* and family Lachnospiraceae.

Although the amino acid sources showed no differences in the overall structure of the final microbial communities, there were differences in community succession. To enumerate the relationship between communities, we calculated the Bray-Curtis dissimilarity between each day and the previous day for each donor and compared the two amino acid sources (Supplemental Figure 6). Comparing each community to the corresponding sample from the previous day demonstrates that the dissimilarity plateaued between days five and seven, independent of amino acid source. Taken together, these data suggest that mucin can sustain diverse fermenting consortia over sequential dilutions in a manner related to the initial community structure, including overall diversity and composition.

## Discussion

Mucins provide an endogenous nutritional resource in the human gut upon which additional nutrients from the hosts diet are overlaid *in vivo*. As such, mucins are one of the most consistent nutritional resources available to gut microorganisms. Although many individual mucin-degrading bacteria have been identified (Tailford et al. 2015), we wanted to examine mucin degradation as a shared community-level process across different host microbiomes. Given that each human hosts a taxonomically distinct gut microbiome, we used fecal samples from three donors to select for mucin-degrading communities over 10-days of sequential passage and assess community composition and metabolic output using 16S rRNA gene sequencing and SCFA quantification, respectively. We were also interested in determining whether the provision of amino acids or oligopeptides would differentially impact community structure, so replicate communities were developed on either tryptone or a proteinogenic amino acid mixture. At the conclusion of the experiment, we observed three distinct mucin-degrading communities across donors with minor differences within a donor’s inoculum on distinct amino acid source. While similarities in final community composition did exist, notable taxonomic differences across donors included the presence of the mucin-degrading specialist *Akkermansia* in the donor three community and the presence of the sulfate-reducing *Desulfovibrio* in the donor two community. Metabolic output differences were most notable for donor two communities, as they produced significantly less acetate and propionate than the other two communities. These findings reinforce the concept that the gut microbiome is taxonomically distinct yet functionally redundant at broad functional levels. However, while all three communities functionally degrade mucin, the taxonomic differences result in distinct metabolic outputs, suggesting that the mechanisms and efficiencies of mucin degradation across individuals are important for understanding how this community-level process impacts human health.

Synthetic microbial communities simplify the study of interaction networks between two or more microorganisms connected by a common function. The study of these microbial functions and the microorganisms that are involved is typified by two approaches, bottom-up (interaction-focused) or top-down (function-focused). Bottom-up approaches intentionally assemble multi-member communities, often using type materials from disparate sources aiming to identify interaction networks. However, strain-level differences across natural communities may impact overall community structure and function (Wu et al. 2015). In the alternate top-down approach, a naturally existing community is cultivated in a controlled environment that allows for selection of naturally co-occurring and interacting strains. However, it is difficult to separate out the effect of an individual carbon source on a microbial community in studies that use a single-inoculation, top-down approach because the initial number and composition of organisms can stifle any minute changes in the microbial composition (Tuncil et al. 2017; Van Herreweghen et al. 2018; 2021). Furthermore, residual stool carbon may result in continued growth of specific organisms within these experiments (Tuncil et al. 2018). To characterize the effect of available nutrients on gut microbial networks, instead of a single-inoculation approach, a consecutive batch-culture approach is becoming more widely used, as it is individual-specific, does not require specialized equipment, and can address second- and third-order interactions such as microbial cheating and cross-feeding (Yao, Chen, and Lindemann 2020; Cheng et al. 2022; Lindemann 2020). Many of these experiments focus on the shift in response to non-native nutrients rather than the analysis of naturally existing microbial networks. Furthermore, microbial community enrichments with specific assemblages are difficult to store, as each microorganism in the community responds differently to long-term storage conditions, necessitating redevelopment in different labs or across multiple experiments (Biclot et al. 2022). With the differences between donor microbiomes seeding these experiments, it is difficult to maintain effective model systems to understanding ecological assembly (Estrela, Sánchez, and Rebolleda-Gómez 2021). However, by using a top-down batch-culture approach with three compositionally different donors in a defined medium with mucin as the sole carbon source, we are better able to identify shared and donor-specific features of mucin degrading gut communities.

In continuous-flow bioreactors that use top-down approaches, such as SHIME, addition of mucin to an existing community grown on complex carbohydrates causes a demonstrable shift in the community structure, supporting *Akkermansia*, Clostridium cluster XIV, and *Ruminococcus* (Van Herreweghen et al. 2018; 2021). Although the SHIME system used in these studies typically have additional carbon sources and mucin wash-out, some observations are consistent with the data presented here. For example, we demonstrate a similar ability to grow *Akkermansia* in a mucin-rich medium in one of the three communities. Contrastingly, members of the Clostridium cluster XIV and genus *Ruminococcus* represented a very small fraction of the final mucin-degrading consortia across all three donors (0.2-1.5% and 0.05-4%, respectively). The results here might be more similar to top-down approaches that limit the availability of other carbon sources, such as the initial descriptions of M-SHIME that add agar-bound mucin (to represent the mucosal mucin) in addition to free mucin in the medium (to represent the free-floating luminal mucin) (Van den Abbeele et al. 2013). At three days post-inoculation in the free mucin medium, members of Firmicutes compose an average of 64% of the community. Although differences between this study and SHIME-based approaches with respect to minor nutrients, amino acids, and bile salt availability are likely to affect final community composition, we also observe communities for which approximately half (average of 48%) is represented by the phylum Firmicutes.

Notable taxonomic similarities across donors include the prevalence of *Bacteroides* in all three final communities despite not a single ASV from this genus being shared across donors. Although some members of the *Bacteroides* have known capacity for mucin degradation, such as *B. fragilis* and *B. thetaiotaomicron*, most are not specialized on this carbon source and instead target dietary fibers (Salyers et al. 1977; Pudlo et al. 2015). However, the breadth of glycoside hydrolases (GH) harbored by these organisms enables subsistence on mucin in the absence of dietary fibers (Glover, Ticer, and Engevik 2022; Sonnenburg et al. 2005) or during strong competition by other community members (Sonnenburg, Chen, and Gordon 2006). Within the *Bacteroides* populations we note two interesting observations across donors; one, that each donor’s microbiome contained an ASV assigned as *B. vulgatus* that had an initial bloom and then disappeared from the communities, and two, there were different dominant *Bacteroides* ASVs by day 5 that were maintained through the remainder of the experiment. *B. vulgatus* is not a known mucin degrader and likely requires other members of the community to liberate sugars from complex substrates; for example, co-cultures with *B. ovatus* on inulin have demonstrated growth dependency of *B. vulgatus* on *B. ovatus*, likely mediated through extracellular GHs (Rakoff-Nahoum, Foster, and Comstock 2016). The other *Bacteroides* that dominated donor communities one and two after day 5 could not be classified beyond genus with the SILVA database. However, manually using NCBI BLAST to assign and confirm taxonomy of these sequences (Supplemental Table 2), we found that they closely matched *B. caccae* (community one), *B. thetaiotaomicron* (community two), and *B. fragilis* (community three). In addition to *B. fragilis,* both *B. caccae* and *B. thetaiotaomicron* are known to degrade mucin (Desai et al. 2016), suggesting that each donor has a unique *Bacteroides* profile capable of mucin consumption that would enable survival in the absence of dietary fiber.

After ten consecutive days of growth on mucin, each donor community maintained ∼30% of its initial diversity, demonstrating that a complex glycoprotein like mucin can sustain relatively diverse microbial communities. While some members of these communities are certainly primary mucin degraders, many of the taxa observed are not. Instead, diversity is at least partially maintained by generalist bacteria feeding on mucin sugars liberated by primary degraders and by bacteria cross-feeding on fermentation end products (Koropatkin, Cameron, and Martens 2012). For example, strains of *B. thetaiotaomicron* are able to liberate sialic acid from mucins but unable to import and catabolize it for carbon and energy (Marcobal et al. 2011). Similarly, *B. bifidum* releases mucin monosaccharides that support other members of the gut microbiome (Bunesova, Lacroix, and Schwab 2018). *Faecalibacterium*, which is a genus of Firmicutes observed in all final donor communities here, is one example of an organism able to use mucin sugars (i.e. GlcNAc) but not intact mucin oligosaccharides (Lopez-Siles et al. 2012). In addition to the liberated monosaccharides, other bacteria could be sustained by fermentation end products. For example, acetate produced by *A. muciniphila* during mucin fermentation can support the growth of the butyrogenic *Eubacterium hallii in vitro* (Belzer et al. 2017). How prevalent this type of cross-feeding across the three donor communities is unknown but could play a significant role in the final amount of SCFAs observed here. Although it is difficult to do more than hypothesize about these metabolic interactions based on 16S rRNA gene sequencing alone, these types of syntrophic interactions are important drivers of overall microbiome function and gut health. Future work will look more directly at these interactions using these communities and co-culture experiments.

Beyond sugar liberation, primary mucin degraders could also sustain other functional groups of bacteria, including sulfate reducers, as suggested by a bloom of *Desulfovibrio* sp. in the donor two community. Many mucin oligosaccharides are capped with sulfate groups that must be removed using sulfatase enzymes before accessing the underlying sugar moieties (Katoh et al. 2017; Luis et al. 2021). Although we do not know the functional capabilities of the strain observed here, other *Desulfovibrio* spp. do not produce sulfatase enzymes capable of this activity (Rey et al. 2013). This suggests that other members of the community must liberate sulfate to sustain the *Desulfovibrio* bloom observed in the donor 2 community. For example, several species belonging to the Bacteroides are capable of removing sulfate groups from mucin (Luis et al. 2021; Ulmer et al. 2014). We propose that the *Desulfovibrio* in community two reduces liberated sulfate into hydrogen sulfide, as has been previously demonstrated in co-culture experiments (Rey et al. 2013), where in the absence of this organism in donor one and three communities, we expect a buildup of sulfate in the culture medium. Where residual sulfate in the colon is likely excreted and is unlikely to affect the host (Florin et al. 1991), high levels of hydrogen sulfide have strong links to colorectal cancer (Wolf et al. 2022). Our study points to the importance of future work aimed at understanding the full metabolic profile of mucin-degrading communities.

Although there were similar trends in community composition across all three donors, for example, of particular interest was the community containing the known mucin-degrading specialist, *Akkermansia*. *Akkermansia* are widely regarded as beneficial members of the gut microbiome and are known to specialize on mucin glycoproteins because of auxotrophies for both GlcNAc and threonine, which are both prevalent in mucin (van der Ark et al. 2018). An initial bloom of *Akkermansia* was observed to represent approximately 20% of the community by day 5, which then dwindled by day 10. This community also contained an ASV assigned as *B. fragilis*, another well-established mucin degrader that increased in abundance as *Akkermansia* decreased, suggestive of competition (Roberton and Stanley 1982; Huang, Lee, and Mazmanian 2011). This apparent competition between *Akkermansia* and *B. fragilis* may be strain- or condition-dependent. Recently, the genus *Akkermansia* has been classified into four species-level phylogroups (AmI-AmIV) based on phylogenomic analyses (Guo et al. 2017; Kirmiz et al. 2020). These genotypic differences are recapitulated in phenotypes, for example, where specific phylogroups of *Akkermansia* synthesize vitamin B_12,_ potentially outcompeting other organisms in a vitamin B_12_-deficient environment (Kirmiz et al. 2020). Other differences may be directly related to mucin metabolism, where phylogroups AmII, AmIII and AmIV retain a larger number of CAZymes, than AmI, which differ in their abundance pattern with phylogroups (Luna et al. 2022; Becken et al. 2021). Although the genomic patterns differ across *Akkermansia*, these differences are unidentifiable within the V4 region of the 16S rRNA gene studied here. Furthermore, strain-level diversity within *B. fragilis*, have identified repeated mutations across individuals in polysaccharide utilizing loci, suggesting continued within-person evolution, which may be directed by competition by other members of the microbiome, such as other members of the *Bacteroides* or, in the case of mucin, *Akkermansia* (Zhao et al. 2019).

Each donor community was developed along triplicate lineages with very little variability in taxonomy, diversity, and composition across replicates throughout the experiment. We interpret these observations to suggest that mucin as the sole carbon source is a strong selective force driving community assembly. These findings are in line with other sequential cultivation experiments on complex polysaccharide structures in which diverse and replicable communities have been developed on mucins (Lou et al. 2023) and dietary fibers (Yao, Chen, and Lindemann 2020). The complexity in glycosidic bonds, which sustain greater species diversity (Midani and David 2023), are likely to drive this selection and limit the ability for any one organism to dominate. In our experiments, the mucin solution is first dialyzed to remove small oligosaccharides (<12kDa) and then filtered (<0.2 uM), resulting in large, soluble glycoproteins that are likely processed extracellularly before transport. This mucin pre-processing step is likely effective, as simple sugars, when present in these types of cultivation experiments can result in communities that are dominated by one or two species (Lou et al. 2023; Yao, Chen, and Lindemann 2020). It will be interesting, however, to determine the inflection point of polymer complexity at which the transition from single-organism culture to multi-species consortium occurs.

In addition to nutrient availability, the absence of specific nutrients in these systems may be influential on selection and community assembly. Auxotrophies, both for vitamins and amino acids, are highly prevalent across many microbial communities (Yu et al. 2022). In similar sequential batch-culture cultivation experiments, the relief of these auxotrophies has a demonstrable effect on final community composition (Yao, Chen, and Lindemann 2020). In the experiments here, common vitamin auxotrophies were alleviated through addition of the ATCC vitamin supplement. However, in hand with amino acid auxotrophies, the abilities of an organism to scavenge amino acids into its cells vary; further, some organisms produce exopeptidases capable of breaking down complex proteins into smaller fragments, as dipeptides or free amino acids, which are then transported into the cell (Zhang et al. 2022). Although the mucin backbone does provide a potential source of amino acids, not all bacteria produce peptidases able to cleave the mucin backbone. Therefore, to ameliorate any amino acid auxotrophy without providing additional carbon sources, we provided a low concentration of free amino acids or tryptone in the fermentation medium and assessed whether the complexity of amino acids influenced the final microbial community structure. The lack of ASVs discriminated by LEfSe analysis comparing the final communities cultivated with the two amino acid sources is in line with observations of bacterial growth in minimal amino acid-free media (Price et al. 2018) and cross-feeding interactions in complex systems (Du et al. 2022). These dynamic interactions by other members of the microbial community may be species- or strain-specific (Ashniev et al. 2022), which argues for a comprehensive analysis of the genomic capacity of these organisms using metagenomics and assembly of species’ genomes to fully determine whether organisms with auxotrophies were maintained in these experiments.

## In summary

Identification of individual mucin-degrading consortia necessitated the use of *in vitro* models, in which mucin was the sole carbon source. For example, in *in vivo* studies in both animals and humans, the nutritional role of mucins is obscured by dietary conditions. Therefore, the batch-model system used in this research provides essential advantages to identify the ecological networks of mucin degradation. However, variability in mucin structure and composition along the length of the intestinal tract (Robbe et al. 2004), secretion of antimicrobial peptides (Bevins 2007), and spatial organization of individual microbes (Mark Welch et al. 2017) are remaining aspects of mucin degradation that are host-relevant and difficult to replicate *in vitro*. Development of these communities *in vitro* using a top-down approach allows the examination of microbe-microbe interactions while accounting for naturally occurring strain heterogeneity amongst core bacterial groups, an advantage not afforded by bottom-up approaches that rely on assembling communities using type strains originating from different sources.

## Data Availability

All raw sequencing data in FASTQ format are available in the NCBI Sequence Read Archive (SRA) BioProject database under accession number PRJNA956585 as BioSamples SAMN34224450-SAMN34224644

## Author Note

We have no known conflict of interest to disclose. This research has complied with all institutional and federal policies regarding the use of human subjects.

## Acknowledgements

We thank anonymous donors 1, 2, and 3, for their generous donation of fecal material and Arno Henzi and Estefani Luna for helping prepare and run 16S rRNA amplicon sequence libraries.

## We declare no conflicts of interest

A.D.F. conceived the project, conducted fermentation experiments, interpreted data, performed statistical analysis, and wrote the manuscript. T.Y. conducted SCFA experiments, interpreted the data, and helped write the paper. S.R.L helped write the paper. G.E.F. helped interpret data and write the paper.

---

Supplemental Figure 1. With the exception of Donor 2 acid production, the optical density (OD, A) and acid production (pH, B) of mucin degrading consortia from the same donor grown with different amino acid sources were consistent. Averages of three lineages for each final donor community (Donor 1, D1; Donor2, D2; Donor3, D3) are displayed with error bars representing standard deviation as calculated in R. Statistically significant differences are calculated by Tukey’s multiple comparisons test with P<0.05. Symbol style: nonsignificant (ns), 0.05 (*), 0.01 (**), 0.001(***) and <0.0001(****)

Supplemental Figure 2. Mucin degrading consortia from donors one and three produced the higher concentration of Isobutyrate (A) and Isovalerate (B) than the the donor two consortium, irrespective of amino acid source. Mean, first and third quartiles for each final donor community (Donor 1, D1; Donor2, D2; Donor3, D3) are represented. Statistically significant differences are calculated by Tukey’s multiple comparisons test. Symbol style: nonsignificant (ns), 0.05 (*), 0.01 (**), 0.001(***) and <0.0001(****)

Supplemental Figure 3. Percent loss of microbial members was approximately equal across donor samples, irrespective of amino acid source. The mean, first and third quartiles for each donor (Donor 1, D1; Donor2, D2; Donor3, D3) of the percent of ASVs as compared to the number of ASVs in the fecal inoculum (day zero) when supplemented with amino acids provided as an equimolar mix (Amino Acids, blue) or peptides (Tryptone, pink) are shown.

Supplemental Figure 4. Mucin sustains diverse fermenting consortia over sequential dilutions that is donor dependent. Average (n=3) relative abundances of top 50 ASVs over 10 days with supplementation of either Amino Acids or Tryptone are shown. Each shade represents a distinct ASV, with the color representing a Family, separated by Bacillota (A), Bacteroidota (B), Desulfobacterota (C), Proteobacteria (D), and Verrucomicrobiota (E).

Supplemental Figure 5. Final consortia had multiple members unique to each donor. Linear discriminant analysis differentiating the combined day 10 communities from both amino acid sources across the donors. Taxa with Linear Discriminant Analysis (LDA) scores >3.0 are shown for ASVs (A) and genera (B).

Supplemental Figure 6. Community structure plateaued between days five and seven, independent of amino acid source. Bray Curtis dissimilarity was calculated between each sample and the corresponding sample from the previous passage along each lineage. Smoothed conditional means calculated for cultures grown with Amino Acids (grey) or Tryptone (orange).

Supplemental Table 1. Microbial community composition was not affected by sequencing run but was different across three donors after 10 days of sequential passage. Beta diversity between selected samples across two sequencing runs and between the final consortium across three donors was compared by ADONIS (PERMANOVA). Abbreviations: df, degrees of freedom; SS, sum of squares. Symbol style: nonsignificant (ns) and 0.001 (**).

Supplemental Table 2. NCBI BLAST confirmed and assigned taxonomy of sequences belonging to the phylum Bacteroidota. Top hit for NCBI classification was selected and compared to the Qiime annotation. ASVs that dominated each community are highlighted in bold with an asterisk adjacent to the NCBI match.

