## Supplemental Figures for "Interindividual Diversity of Human Gut Mucin-Degrading Microbial Consortia"

### Slide 1
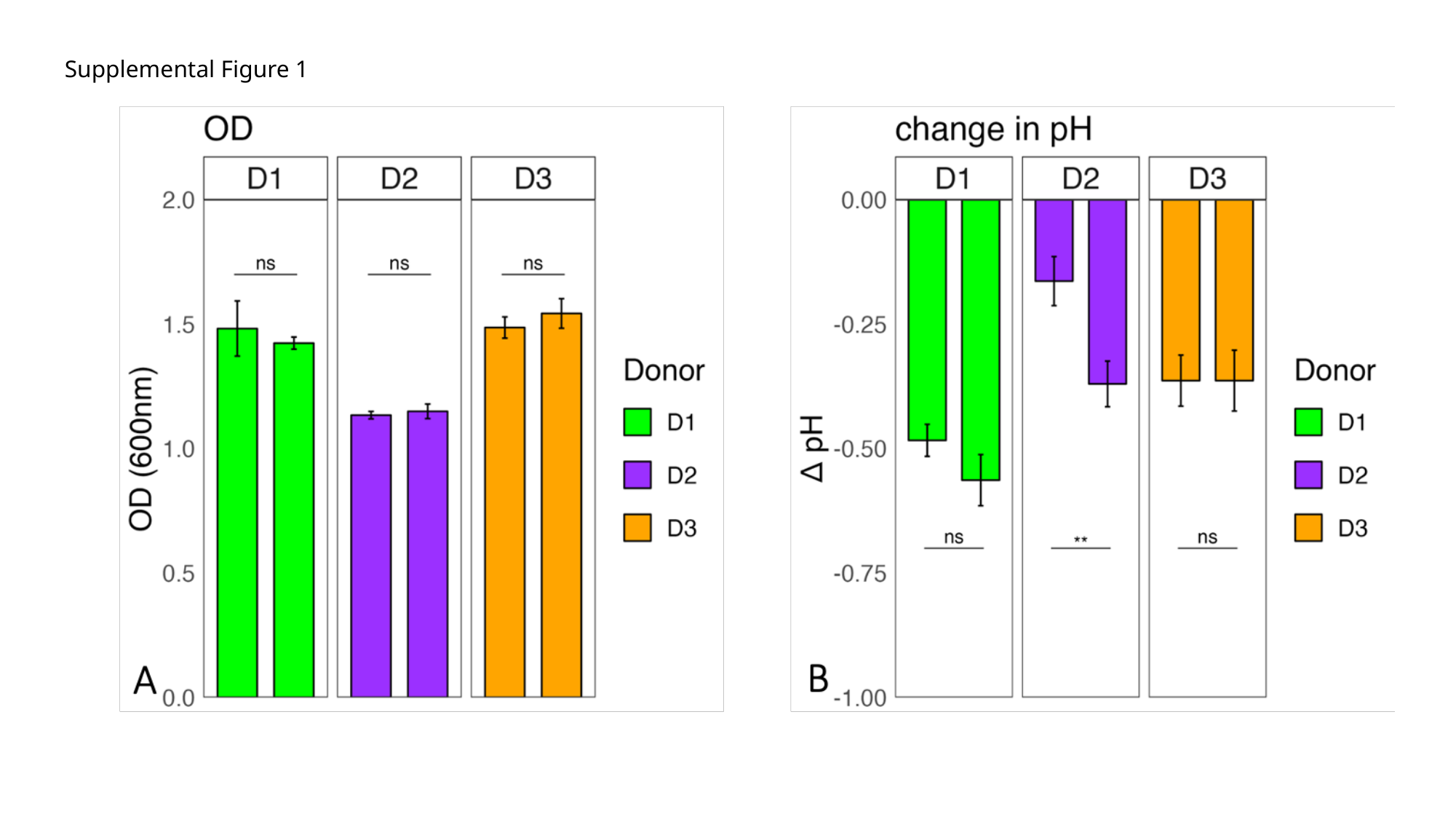

#### Supplemental Figure 1

### Slide 2
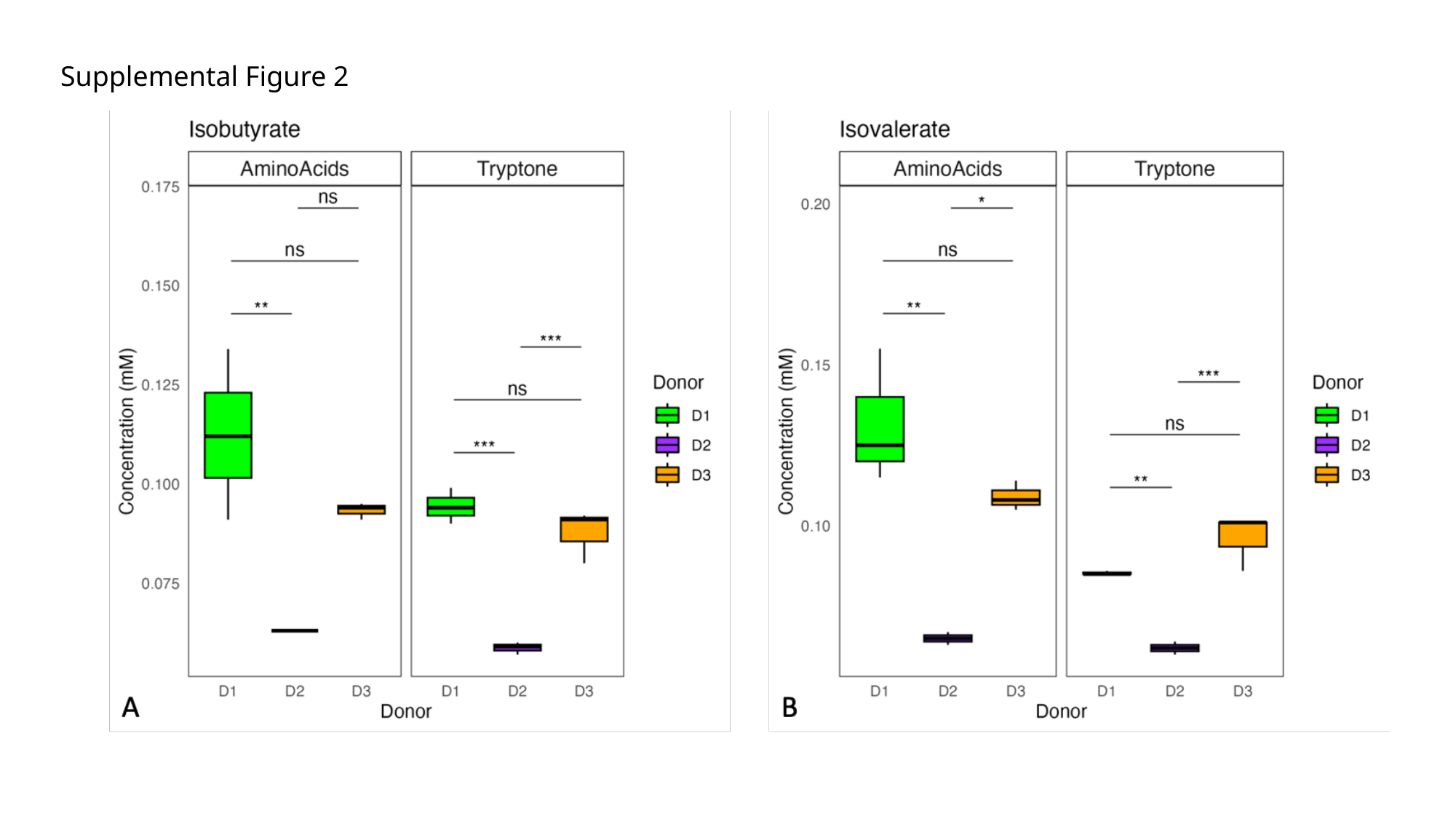

Supplemental Figure 2

### Slide 3
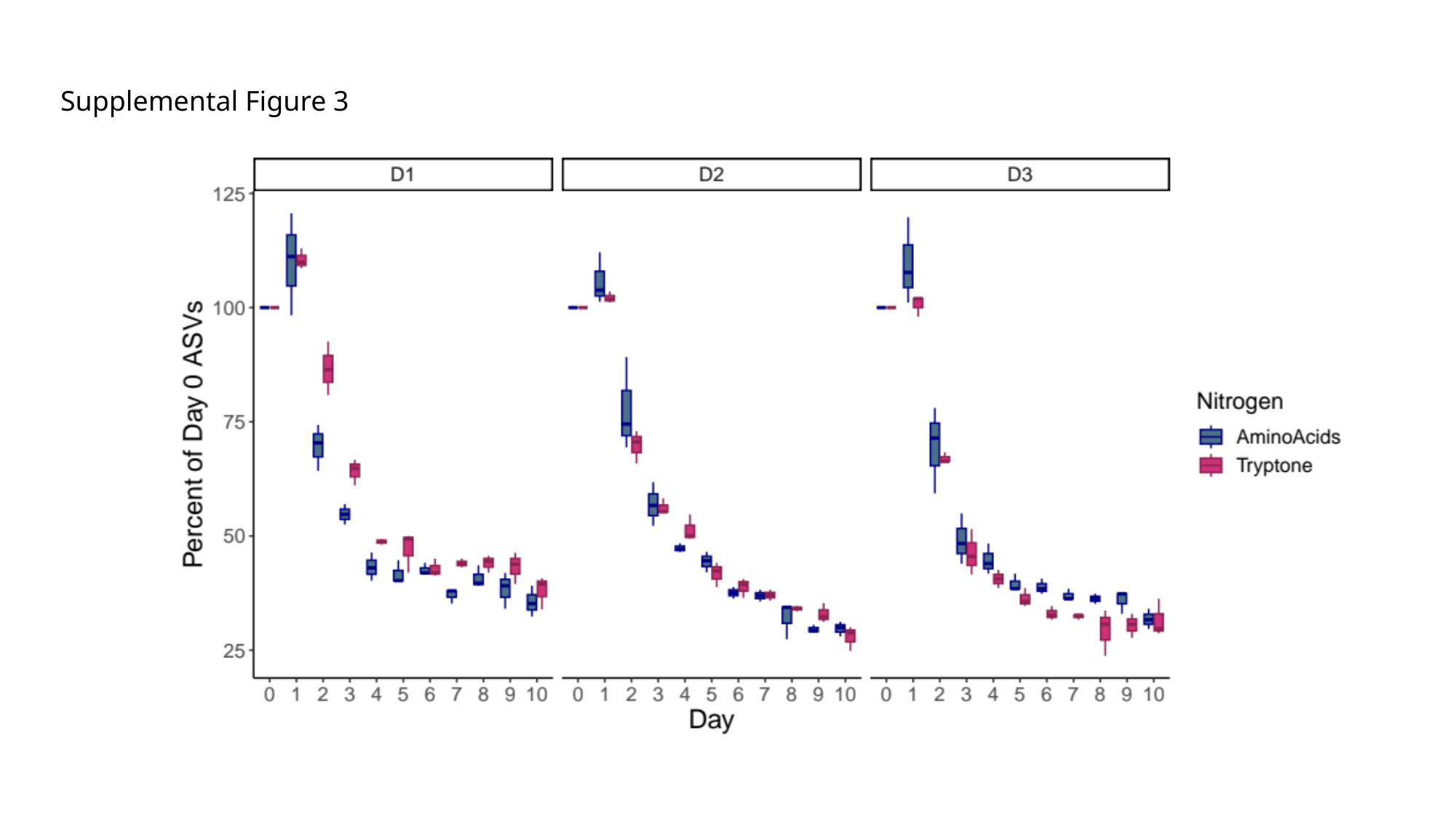

Supplemental Figure 3

### Slide 4
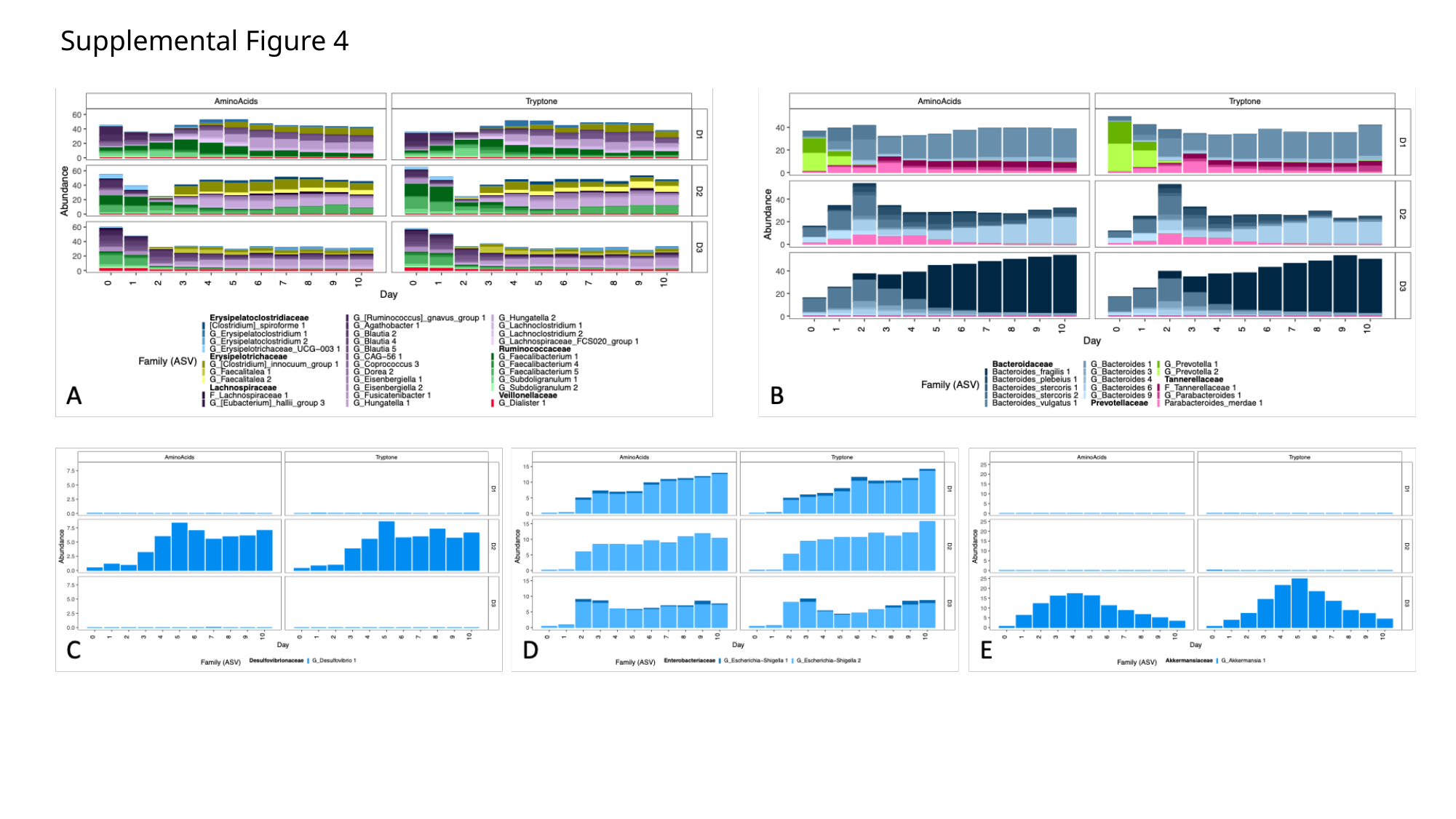

Supplemental Figure 4

### Slide 5
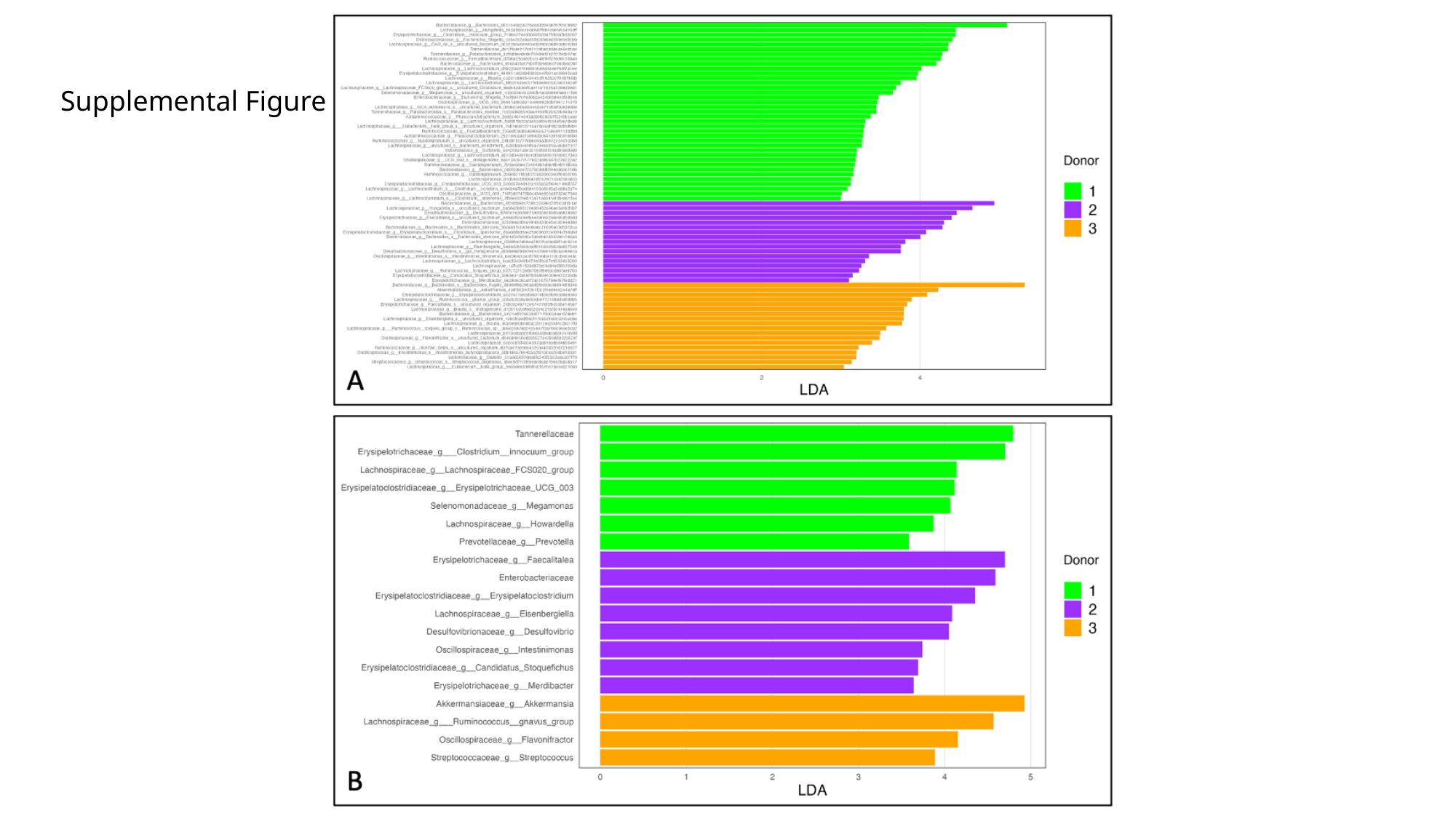

Supplemental Figure 5

### Slide 6
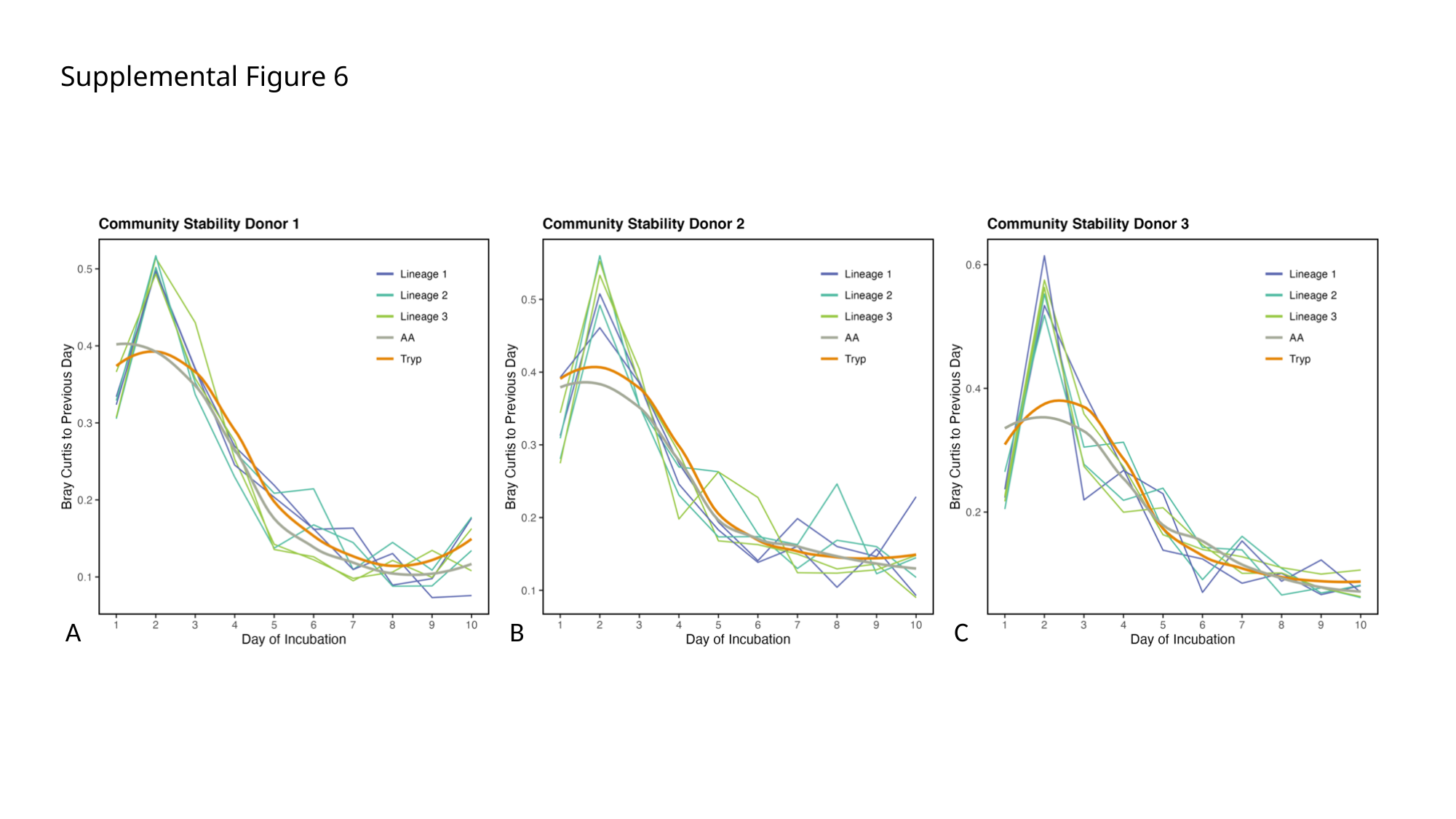

Supplemental Figure 6
A
B
C
