## Supplemental Tables for "Interindividual Diversity of Human Gut Mucin-Degrading Microbial Consortia"

**Supplemental Table 1**

|  |  | ***df*** | ***SS*** | ***R2*** | ***F*** | ***Pr(>F)*** | **Perms** |
| --- | --- | --- | --- | --- | --- | --- | --- |
| Samples sequenced on run 1 and run 2 (n = 6 per group) | Bray Curtis | *1* | *0.010414* | *0.0032369* | *0.0325* | *0.96 (ns)* | 999 |
|  | Unweighted | 1 | 0.083236 | 0.052774 | 0.5571 | 0.633 (ns) | 999 |
|  | Weighted | 1 | 0.0008892 | 0.0025508 | 0.0256 | 0.947 (ns) | 999 |
|  | Jaccard | 1 | 0.15304 | 0.044219 | 0.4627 | 0.848 (ns) | 999 |
| Donor Communities at Day 10  (n = 6 per group) | Bray Curtis | 3 | 3.9001379 | 0.8928368 | 44.43502 | 0.001*** | 999 |
|  | Unweighted | 3 | 1.213736 | 0.7837684 | 19.33158 | 0.001*** | 999 |
|  | Weighted | 3 | 0.39984043 | 0.8531772 | 30.99163 | 0.001*** | 999 |
|  | Jaccard | 3 | 3.9475482 | 0.8191273 | 24.15334 | 0.001*** | 999 |

**Supplemental Table 2**

| ASV | NCBI Match | Qiime Annotation |
| --- | --- | --- |
| 0cc2f79a3e3d31e1d291a8e083e611ab | Bacteroides_finegoldii | c__Bacteroidia o__Bacteroidales f__Bacteroidaceae g__Bacteroides |
| 1b6ffc0a05a7be8d4f4d95425832d107 | Bacteroides_caccae | c__Bacteroidia o__Bacteroidales f__Bacteroidaceae g__Bacteroides s__Bacteroides_caccae |
| 1c3b3d6b5540ae4459f826420b48da10 | Parabacteroides_merdae | c__Bacteroidia o__Bacteroidales f__Tannerellaceae g__Parabacteroides s__Parabacteroides_merdae |
| 1ef97ff90aed6940d9fbd604ce20da5d | Bacteroides_caccae | c__Bacteroidia o__Bacteroidales f__Bacteroidaceae g__Bacteroides s__Bacteroides_caccae |
| 20cf13f228f8d16b499e9493dd2ee208 | Bacteroides_uniformis | c__Bacteroidia o__Bacteroidales f__Bacteroidaceae g__Bacteroides |
| 2d3f536ce7257608ddf6f44a800c218b | Bacteroides_nordii | c__Bacteroidia o__Bacteroidales f__Bacteroidaceae g__Bacteroides |
| 2ed7801e4cef3b35f4140acd2f1101e7 | Bacteroides_intestinalis | c__Bacteroidia o__Bacteroidales f__Bacteroidaceae g__Bacteroides s__Bacteroides_intestinalis |
| 3a5dc0b707e689a2beeeb1e6c17d9f6c | Bacteroides_clarus | c__Bacteroidia o__Bacteroidales f__Bacteroidaceae g__Bacteroides s__Bacteroides_clarus |
| 3cf6dd6edede75f439d74272791b67ac | Parabacteroides_pekinense | c__Bacteroidia o__Bacteroidales f__Tannerellaceae g__Parabacteroides |
| 4f13b31757426267a1b923846bcfe43d | Bacteroides_uniformis | c__Bacteroidia o__Bacteroidales f__Bacteroidaceae g__Bacteroides s__Bacteroides_uniformis |
| 507d64a7f56fccec9f413747b5842972 | Parabacteroides_distasonis | c__Bacteroidia o__Bacteroidales f__Tannerellaceae g__Parabacteroides |
| 50da935cb4343ba6c21695a03d5353ca | Bacteroides_stercoris | c__Bacteroidia o__Bacteroidales f__Bacteroidaceae g__Bacteroides s__Bacteroides_stercoris |
| 5401edfc1902e0f71700034ae1f78eb1 | Bacteroides_ovatus | c__Bacteroidia o__Bacteroidales f__Bacteroidaceae g__Bacteroides |
| 6501054fb595c538eb4033530612d4a3 | Bacteroides_stercoris | c__Bacteroidia o__Bacteroidales f__Bacteroidaceae g__Bacteroides s__Bacteroides_stercoris |
| 6521f34e85eb9034e3bd9d4e58dff50a | Bacteroides_ovatus ; Bacteroides_xylanisolvens | c__Bacteroidia o__Bacteroidales f__Bacteroidaceae g__Bacteroides |
| 6bd982a66b0c74116b5c1cd9b824c113 | Bacteroides_cellulosyliticus | c__Bacteroidia o__Bacteroidales f__Bacteroidaceae g__Bacteroides |
| 6e14788f3f3cb78ee7fa0348a0c2cf46 | Bacteroides_plebeius | c__Bacteroidia o__Bacteroidales f__Bacteroidaceae g__Bacteroides s__Bacteroides_plebeius |
| 7b7f3706e2803ced3a3eddfa7887e7dc | Bacteroides_sartorii | c__Bacteroidia o__Bacteroidales f__Bacteroidaceae g__Bacteroides s__Bacteroides_sartorii |
| 84618cc630790f97e68fed6ac3a55c68 | Bacteroides_coprocola | c__Bacteroidia o__Bacteroidales f__Bacteroidaceae g__Bacteroides s__Bacteroides_coprocola |
| 8974faa17be540e9a3a18be3919846bb | Bacteroides_thetaiotaomicron | c__Bacteroidia o__Bacteroidales f__Bacteroidaceae g__Bacteroides |
| 8f2ddb8d1729fc35598d70fbcbd1b1af | Bacteroides_thetaiotaomicron* | c__Bacteroidia o__Bacteroidales f__Bacteroidaceae g__Bacteroides |
| 97d34dbaf75b1ffbb56b6d7b8db933bf | Bacteroides_ovatus | c__Bacteroidia o__Bacteroidales f__Bacteroidaceae g__Bacteroides |
| b0c83858accb36fae1309230e71f31a3 | Bacteroides_coprocola | c__Bacteroidia o__Bacteroidales f__Bacteroidaceae g__Bacteroides s__Bacteroides_coprocola |
| b57b0a37bd78a259c27a9cc823483311 | Bacteroides_vulgatus | c__Bacteroidia o__Bacteroidales f__Bacteroidaceae g__Bacteroides s__Bacteroides_vulgatus |
| b5bf467bb53eb088c18e684d8656f904 | Bacteroides_clarus | c__Bacteroidia o__Bacteroidales f__Bacteroidaceae g__Bacteroides s__Bacteroides_clarus |
| bd125abcbb6d6ed03a18c0515fee1db4 | Parabacteroides_distasonis | c__Bacteroidia o__Bacteroidales f__Tannerellaceae g__Parabacteroides |
| c06b712348f8542720421722120be38e | Bacteroides_uniformis | c__Bacteroidia o__Bacteroidales f__Bacteroidaceae g__Bacteroides |
| ca1392eaa969b2980a69cc34a327e112 | Bacteroides_dorei | c__Bacteroidia o__Bacteroidales f__Bacteroidaceae g__Bacteroides s__Bacteroides_vulgatus |
| cece12865e81c96a523b3bb6a4511561 | Bacteroides_vulgatus | c__Bacteroidia o__Bacteroidales f__Bacteroidaceae g__Bacteroides s__Bacteroides_vulgatus |
| d051e46a2ac73aead2b4a876701c9862 | Bacteroides_caccae* | c__Bacteroidia o__Bacteroidales f__Bacteroidaceae g__Bacteroides |
| d0a48a8fafdce31a7583434c537587c2 | Parabacteroides_distasonis | c__Bacteroidia o__Bacteroidales f__Tannerellaceae g__Parabacteroides |
| d8999f85296a9d9f0eb3a48834df9d46 | Bacteroides_fragilis* | c__Bacteroidia o__Bacteroidales f__Bacteroidaceae g__Bacteroides s__Bacteroides_fragilis |
| db79739512d1a8e2ea6457e99ff96b8f | Parabacteroides_distasonis | c__Bacteroidia o__Bacteroidales f__Tannerellaceae g__Parabacteroides s__Parabacteroides_distasonis |
| e1dd787a45b6582d7dc397295640fed0 | Bacteroides_clarus | c__Bacteroidia o__Bacteroidales f__Bacteroidaceae g__Bacteroides s__Bacteroides_clarus |
| f00b40d9dfd549fc8240a0dec837bf53 | Bacteroides_sartorii | c__Bacteroidia o__Bacteroidales f__Bacteroidaceae g__Bacteroides s__Bacteroides_sartorii |
